# DENR/MCTS1 knockdown modulates repeat-associated non-AUG translation

**DOI:** 10.1101/2021.08.27.457993

**Authors:** Katelyn M. Green, Shannon L. Miller, Indranil Malik, Peter K. Todd

**Affiliations:** Department of Neurology, University of Michigan; Cellular and Molecular Biology Graduate Program, University of Michigan; Department of Chemistry, Department of Human Genetics, and Center for RNA Biomedicine, University of Michigan; VA Ann Arbor Healthcare System, Ann Arbor, MI

**Keywords:** RAN translation, translation initiation, C9orf72, ALS, FXTAS, eIF2A, DENR, MCTS1, eIF2D

## Abstract

Repeat associated non-AUG (RAN) translation of mRNAs containing repeat-expansion mutations produces toxic peptides in neurons of patients suffering from neurodegenerative diseases. Recent findings indicate that RAN translation in diverse model systems is not inhibited by cellular stressors that impair global translation through phosphorylation of the alpha subunit of eIF2, the essential eukaryotic translation initiation factor that brings the initiator tRNA to the 40S ribosome. Using *in vitro*, cell-based, and *Drosophila* models, we examined the role of alternative ternary complex factors that may function in place of eIF2, including eIF2A, eIF2D, and DENR/MCTS1. Among these factors, DENR knockdown had the greatest inhibitory effect on RAN translation of expanded GGGGCC and CGG repeat reporters, and its reduction improved survival of *Drosophila* expressing expanded GGGGCC repeats. Taken together, these data support a role for alternative initiation factors in RAN translation and suggest they may serve as novel therapeutic targets in neurodegenerative disease.

## Introduction

While less efficient at initiating translation than canonical AUG start codons, ribosome profiling experiments suggest that non-AUG codons define a significant fraction of all cellular translation initiation sites (1, 2). In particular, upstream open reading frames (uORFs) are enriched for non-AUG start codons (1-5), and proteins synthesized with non-AUG start codons often play important roles related to the cellular stress response (reviewed in (6)). As such, non-AUG initiation event can be subject to different regulatory mechanisms than canonical AUG-initiated translation during the integrated stress response (ISR). The ISR reduces global cellular translation by inducing phosphorylation of the essential eukaryotic initiation factor eIF2 at serine 51 of its alpha subunit. Phosphorylation prevents eIF2 from exchanging GDP for GTP, which subsequently prevents eIF2 from binding and delivering the initiator methionine tRNA (Met-tRNA_i_^Met^) to the 40S ribosome. While this strategy is effective at dramatically decreasing most AUG-initiated translation, some non-AUG initiation events are spared or upregulated during the ISR (2, 6, 7).

One model for how specific non-AUG initiation events evade downregulation during ISR activation depends on the ability of specific non-AUG start codons to receive initiator tRNAs independent of eIF2. eIF2A, a non-essential, monomeric protein, can deliver initiator tRNAs to the pre-initiation ribosome and has been shown to act in place of eIF2 during the ISR (7-11). It represents an intriguing therapeutic target, as mice lacking eIF2A survive to adulthood with no gross abnormalities (12). Additionally, eIF2D and DENR/MCTS1 are factors that normally promote ribosome recycling and translation re-initiation (13-15), but can also function in place of eIF2 to promote translation initiation and are not inhibited by ISR activation (16, 17). In addition to binding and delivering the Met-tRNA_i_^Met^, each of these factors can also deliver non-methionyl tRNAs cognate to near-AUG codons used for initiation (8, 11, 16).

While non-AUG initiation is important in normal cellular processes, its mis-regulation has also been implicated in human disease (3, 6, 18). Repeat expansion mutations associated with several neurodegenerative and neuromuscular diseases undergo a process known as repeat-associated non-AUG (RAN) translation (18-22). RAN translation initiates upstream of or within the expanded repeats, and results in translation through the repetitive RNA sequence. This produces RAN peptides that contain large, repetitive amino acid sequences that are often aggregation-prone and neurotoxic in model systems.

By definition, RAN translation exclusively utilizes non-AUG start codons for synthesis of these toxic proteins. Recently, our lab and others showed that RAN translation of expanded GGGGCC repeats associated with C9orf72 amyotrophic lateral sclerosis (ALS) and frontotemporal dementia (FTD, “C9RAN”), as well as of expanded CGG repeats associated with fragile X-associated tremor/ataxia syndrome (FXTAS, “CGG RAN”), behave like some other non-AUG translation events and is increased following ISR activation, while global translation is simultaneously downregulated (23-26). Here, we investigate the ability of C9 and CGG RAN translation to utilize the eIF2-alternatives eIF2A, eIF2D, and DENR/MCTS1 during translation initiation. We find evidence that DENR/MCTS1 and eIF2A support RAN translation of both CGG and GGGGCC repeats under specific conditions. Knocking down DENR, and to a lesser degree eIF2A, modestly improves repeat-mediated toxicity in *Drosophila* expressing GGGGCCx28 repeats. These data suggest a role for alternative ternary complex factors in RAN translation.

## Results

### Loss of eIF2A reduces C9 and CGG RAN translation in vitro

We first assessed a role for eIF2A in C9 or CGG RAN translation by expressing nanoluciferase (NLuc) reporter mRNAs containing either 70 GGGGCC in the glycine-alanine (GA) reading frame or 100 CGG repeats in the +1, poly-glycine reading frame (23, 27) in *in vitro* translation lysates generated from wildtype (WT) and CRISPR-mediated eIF2A knockout (KO) HAP1 cells (**Fig. 1A-B**) (28, 29). Relative to WT lysates consisting of the same total protein concentration, basal translation levels of a canonically translated AUG-NLuc mRNA were significantly reduced in eIF2A KO lysates (**Fig. 1C**). However, after adjusting for the basal difference, RAN translation was significantly reduced in the absence of eIF2A relative to expression of the canonical AUG-NLuc reporter mRNA (**Fig. 1D**). This was true for both the polyGA C9RAN and the +1 CGG RAN translation reporter mRNAs (**Fig. 1D**), and is consistent with a previous report that loss of eIF2A reduces polyGA production in multiple cells lines and chick embryos (25) as well as RAN translation from CCUG/CAGG repeats associated with myotonic dystrophy type 2 (30).

**Figure 1:**
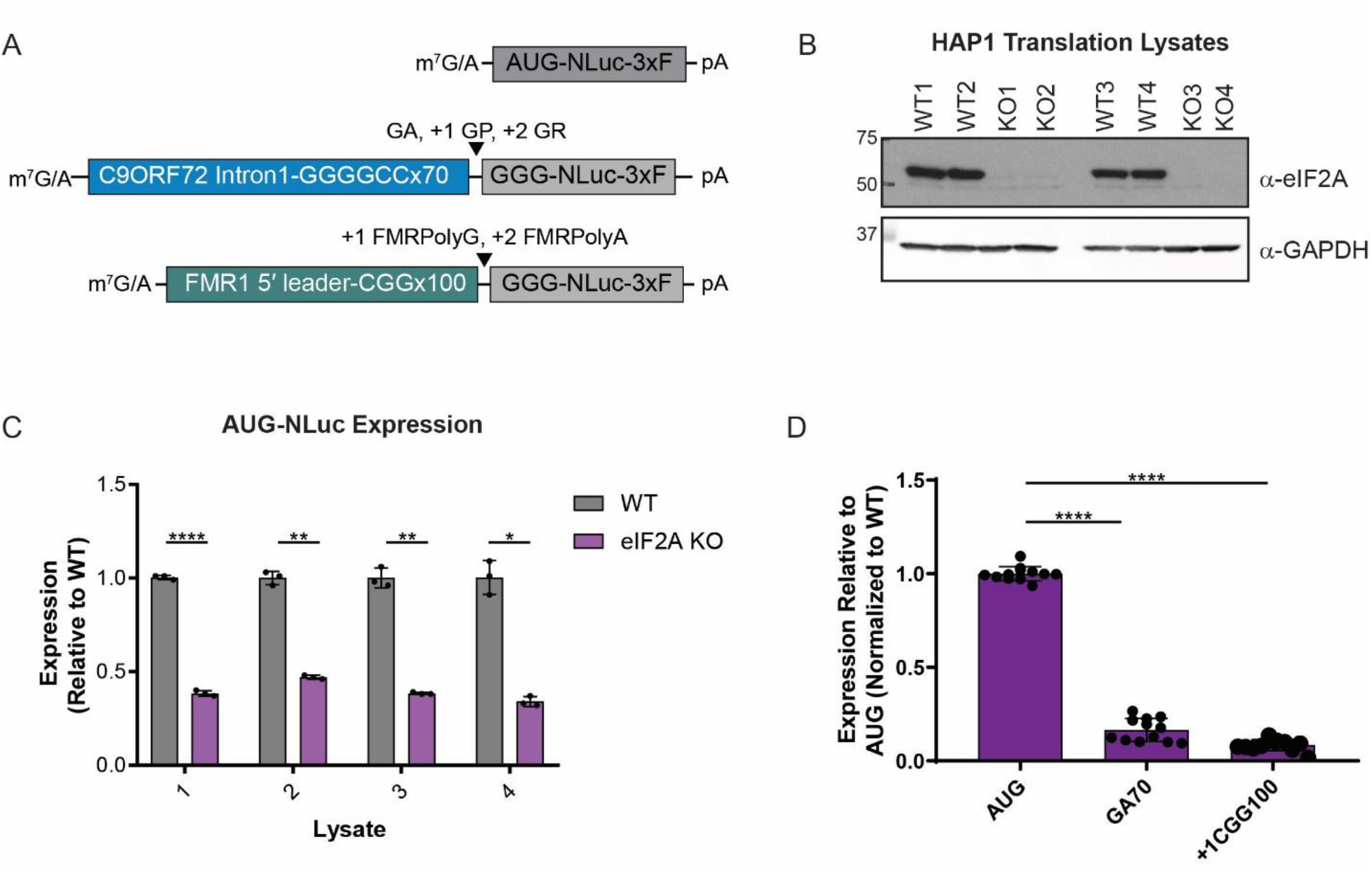
RAN translation is suppressed *in vitro* in eIF2A KO cell lysates. (A) Schematic of Nanoluciferase (NLuc) reporter mRNAs used in rabbit reticulocyte lysate (RRL) *in vitro* translation assays. 3xF = 3x Flag tag, GA = glycine-alanine, GP = glycine-proline, GR = glycine-arginine, +1 FMRpolyG= the glycine reading frame product of the CGG repeat ; +2 FMRpolyA= the alanine reading frame product of the CGG repeat. (B) Western blot of wildtype (WT) and eIF2A knockout (KO) lysates probed with an anti-eIF2A antibody to confirm loss of eIF2A expression. GAPDH is used as a loading control. (C) NLuc expression of AUG-NLuc reporter mRNA in four independently generated sets of WT and KO translation lysates with expression normalized to the WT lysate in each set, n=3 (D) Expression of AUG-NLuc, GA70-NLuc, and +1CGG100-NLuc reporter mRNAs in eIF2A KO lysates, relative to WT lysates, after controlling for the difference in AUG-NLuc expression between paired lysates, n=12. (C) Two-way ANOVA with Sidak’s multiple comparison test. (D) One-way ANOVA with Dunnett’s multiple comparison test. * p< 0.05, ** p< 0.01, **** p< 0.0001

However, in HEK293 cells, siRNA-mediated knockdown (KD) of eIF2A had little effect on C9 and CGG RAN translation from NLuc reporter plasmids co-transfected with FFLuc controls (**Fig. 2A-C**, and **Supplemental Fig.1A**). While it modestly decreased polyGA expression 10-20% relative to cells treated with a non-targeting siRNA (**Fig. 2B & D)**, no inhibitory effect was observed for the glycine-proline (GP) and glycine-arginine (GR) frames, nor for the +1 and +2 reading frames for the CGG repeat (**Fig. 2B & C**). Furthermore, no inhibition was observed when mRNA reporters like those used in the *in vitro* lysate experiments were directly transfected into cells with eIF2A KD, or for NLuc reporters that initiate with a CUG or ACG start codon but lack an expanded repeat (**Fig. 2D & E**).

**Figure 2:**
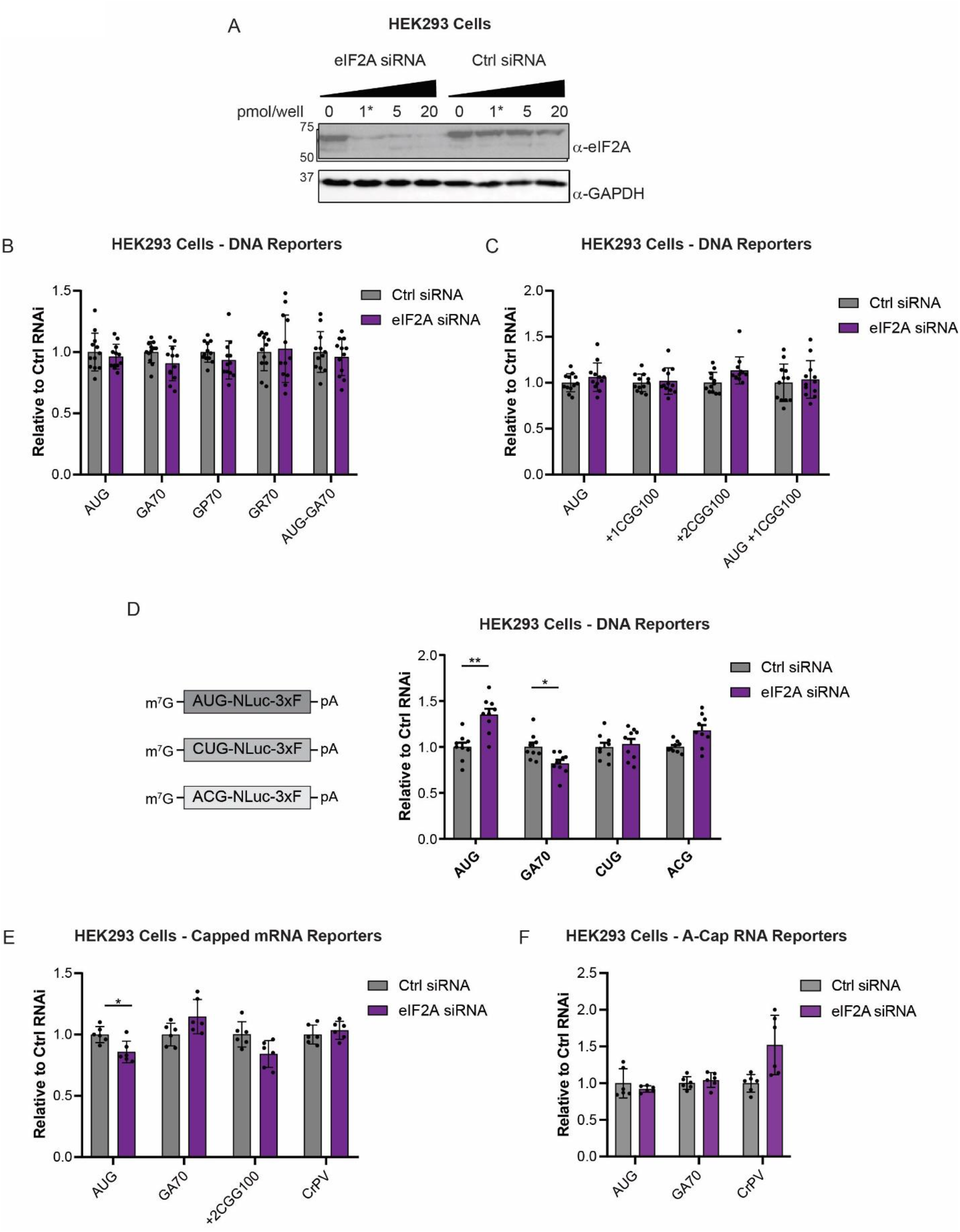
eIF2A knockdown does not alter RAN translation in transfected cells. (A) Western blot showing efficiency of eIF2A KD in HEK293 cells following increasing concentrations of eIF2A siRNA. The starred lane indicates siRNA concentration used in subsequent experiments. GAPDH was used as a loading control. (B-C) NLuc expression of indicated reporters expressed from DNA plasmids in HEK293 cells 24 hours post transfection with non-targeting or eIF2A siRNAs. NLuc levels are expressed relative to levels in cells transfected with the non-targeting siRNA, n=12. (D) Schematic of CUG and ACG-NLuc reporter mRNAs, and NLuc expression of indicated reporters expressed from DNA plasmids transfected into HEK293 cells 24 hours post transfection with non-targeting or eIF2A siRNAs. NLuc levels are expressed relative to levels in cells transfected with the non-targeting siRNA, n=9. (E) NLuc expression of indicated reporters expressed from ARCA-capped mRNAs transfected into HEK293 cells 24 hours post transfection with non-targeting or eIF2A siRNAs. NLuc levels are expressed relative to levels in cells transfected with the non-targeting siRNA, n=6. (F) NLuc expression of indicated reporters expressed from A-capped RNAs transfected into HEK293 cells 24 hours post transfection with non-targeting or eIF2A siRNAs. NLuc levels are expressed relative to levels in cells transfected with the non-targeting siRNA, n=6. All graphs represent mean with errors +/- standard deviation. * p< 0.05, ** p< 0.01, Two-way ANOVA with Sidak’s multiple comparison test.

Previous studies showing inhibition of C9RAN translation in the polyGA frame used bi-cistronic reporters with the GGGGCC repeat placed in the second cistron (25). In this sequence context, cap-independent RAN translation of the GGGGCC repeat could contribute a greater proportion of the luminescent signal than the signal generated from our functionally capped, monocistronic reporter mRNAs. To therefore test if cap-independent translation initiation is more sensitive to eIF2A levels, we next transfected eIF2A KD HEK293 cells with A-capped reporter RNAs (23, 27). Unlike the functional m^7^G, the A-cap cannot interact with the cap-binding factors to support cap-dependent translation initiation. As previously reported (23, 27), relative to functionally capped RAN reporters, the A-capped RAN reporters supported significantly lower levels of translation (**Supplemental Fig. 1B**). However, the remaining fraction of translation observed, which likely occurs through a cap-independent mechanism, also was not inhibited be eIF2A KD (**Fig. 2F**). Consequently, substantial reduction of eIF2A is not sufficient to reduce C9 or CGG RAN translation that occurs through either cap-dependent or cap-independent mechanisms in reporter transfected cells.

### DENR/MCTS1 KD reduces RAN translation in HEK cells

To next determine if either eIF2D and/or DENR/MCTS1 are involved in C9 or CGG RAN translation, we expressed RAN reporters in HEK293 cells following KD of each protein. Reduced levels of eIF2D did not inhibit C9 or CGG RAN translation in any reading frame tested (**Supplemental Fig. 2A-D**). It did, however, modestly reduce expression of the ACG-initiated, no repeat reporter, but had no effect on a no repeat CUG-initiated reporter (**Supplemental Fig. 2E**).

In contrast, reduction of DENR/MCTS1 levels, through use of a DENR-specific siRNA, significantly reduced NLuc expression of C9 and CGG RAN reporters across all reading frames assayed, without significantly altering AUG-NLuc expression (**Fig. 3A-D**). The reduction of polyGA synthesis, but unchanged AUG-NLuc expression following DENR KD was confirmed by Western blot using an antibody against the c-terminal FLAG tag present on each reporter (**Fig. 3B**).

**Figure 3:**
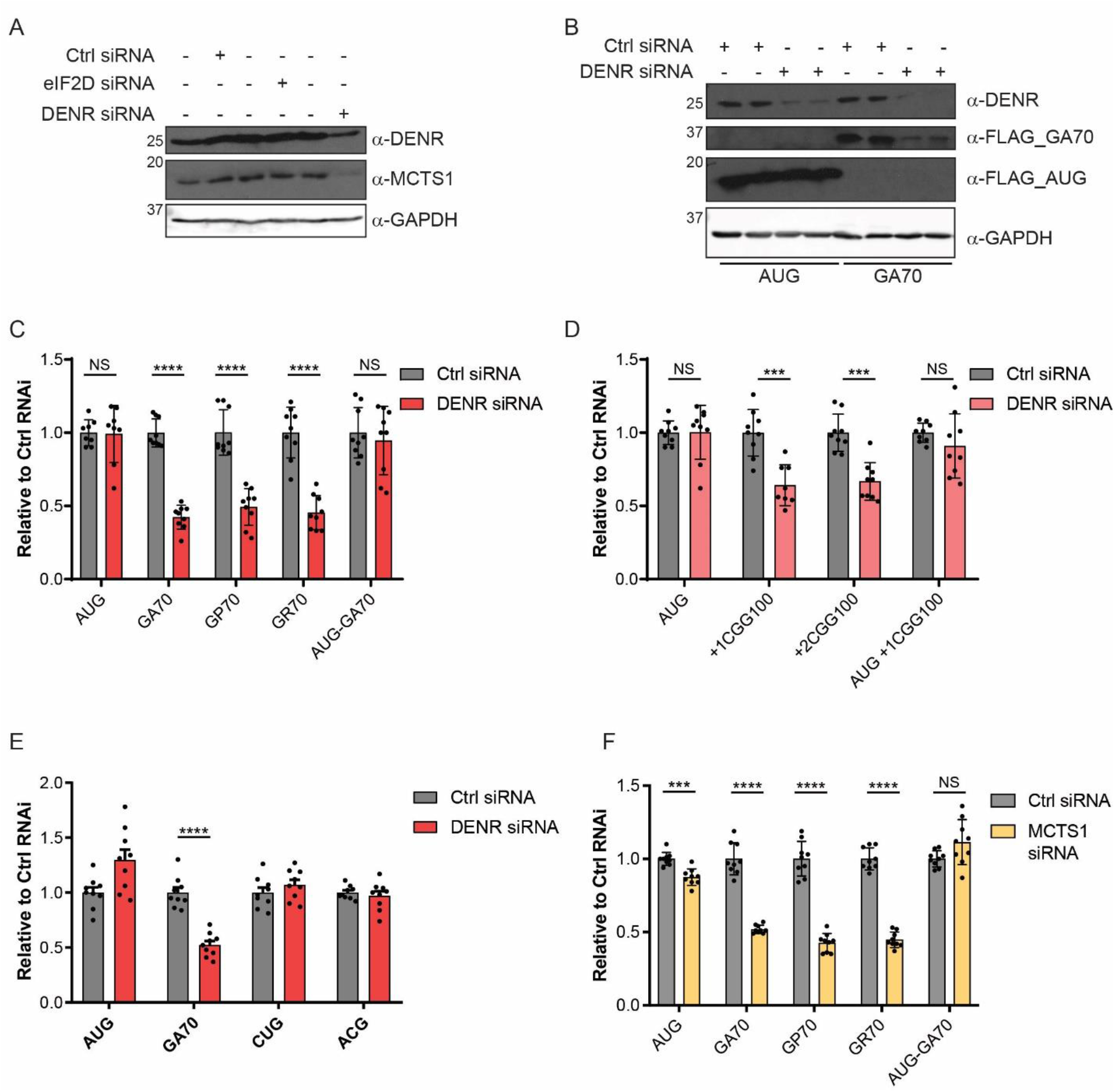
DENR/MCTS1 knockdown selectively suppresses RAN translation. (A) Western blot showing efficiency of DENR and MCTS1 KD in HEK293 cells 48 hours post transfection with an DENR targeting siRNA. GAPDH was used as a loading control. (B) Western blots showing effects of DENR KD on expression of AUG-NLuc and GA70-NLuc reporters expressed in HEK293 cells, 48 hours post KD and 24 hours post reporter transfection. NLuc reporters were probed with an antibody targeting their c-terminal FLAG tag. GAPDH was used as a loading control. (C-E) NLuc expression of indicated reporters expressed from DNA plasmids transfected into HEK293 cells 24 hours post transfection with non-targeting or DENR siRNAs. NLuc levels are expressed relative to levels in cells transfected with the non-targeting siRNA, n=9. (F) NLuc expression of indicated reporters expressed from DNA plasmids in HEK293 cells 24 hours post transfection with non-targeting or MCTS1 siRNAs. NLuc levels are expressed relative to levels in cells transfected with the non-targeting siRNA, n=9. All graphs represent mean with errors +/- standard deviation. *** p< 0.001, **** p< 0.0001, Two-way ANOVA with Sidak’s multiple comparison test.

Interestingly, the effect of DENR KD on GGGGCCx70 and CGGx100 reporter expression was dependent on their non-AUG initiation, as reporters with an AUG start codon inserted upstream of either repeat, to drive canonical translation in the GA or +1 CGG reading frames, were not significantly inhibited (**Fig. 3C & D**). Furthermore, this effect was specific to repeat-associated non-AUG translation, as opposed to no-repeat non-AUG translation, as DENR KD did not reduce expression of CUG-NLuc or ACG-NLuc reporters (**Fig. 3E**).

Although DENR KD had no effect on expression of the highly stable AUG-NLuc protein, it did reduce expression of the less stable, co-transfected AUG-firefly luciferase (FFLuc) reporter, indicating that DENR KD’s inhibitory affect is not specific to RAN translation (**Supplemental Fig. 3A**). However, there was no significant difference in the polysome/monosome ratio in polysome profiles from control vs DENR KD cells, suggesting that DENR KD does not cause global translational inhibition (**Supplemental Fig. 3B**).

DENR functions as a heterodimer with MCTS1, and knockdown of one protein leads to the depletion of the other (**Figure 3A**). To verify the specific effect we observed on C9RAN translation using a DENR-targeting siRNA, we also knocked-down this complex using an MCTS1-targeting siRNA (**Supplemental Fig. 3C**). As expected, MCTS1 KD produced very similar results to DENR KD, having a larger inhibitory effect on C9RAN translation than on translation from either of the AUG-driven NLuc controls (**Figure 3F**), but also significantly reduced expression of the FFLuc transfection control (**Supplemental Fig. 3D**). Together, these data support a role for the DENR/MCTS1 complex in supporting C9 and CGG RAN translation.

### eIF2A, eIF2D, and DENR knockdowns do not prevent increased C9 and CGG RAN translation upon ISR induction

In previous studies, eIF2A was most critical for translation initiation under conditions of cellular stress, when functional eIF2 levels were limited by ISR-mediated phosphorylation of eIF2α at serine 51 (7, 31, 32). We thus specifically assessed the ability of eIF2A, eIF2D, and DENR/MCTS1 KD to regulate C9 and CGG RAN translation following induction of ER stress. Consistent with previous findings (23-26), 2 µM thapsigargin (TG) treatment for five hours selectively increased C9RAN translation expression in HEK293 cells, while inhibiting expression of AUG-initiated control reporters (**Figure 4A to C**). However, KD of eIF2A, eIF2D, and DENR/MCTS1 did not prevent this increase in RAN levels (**Figure 4A to C**).

**Figure 4:**
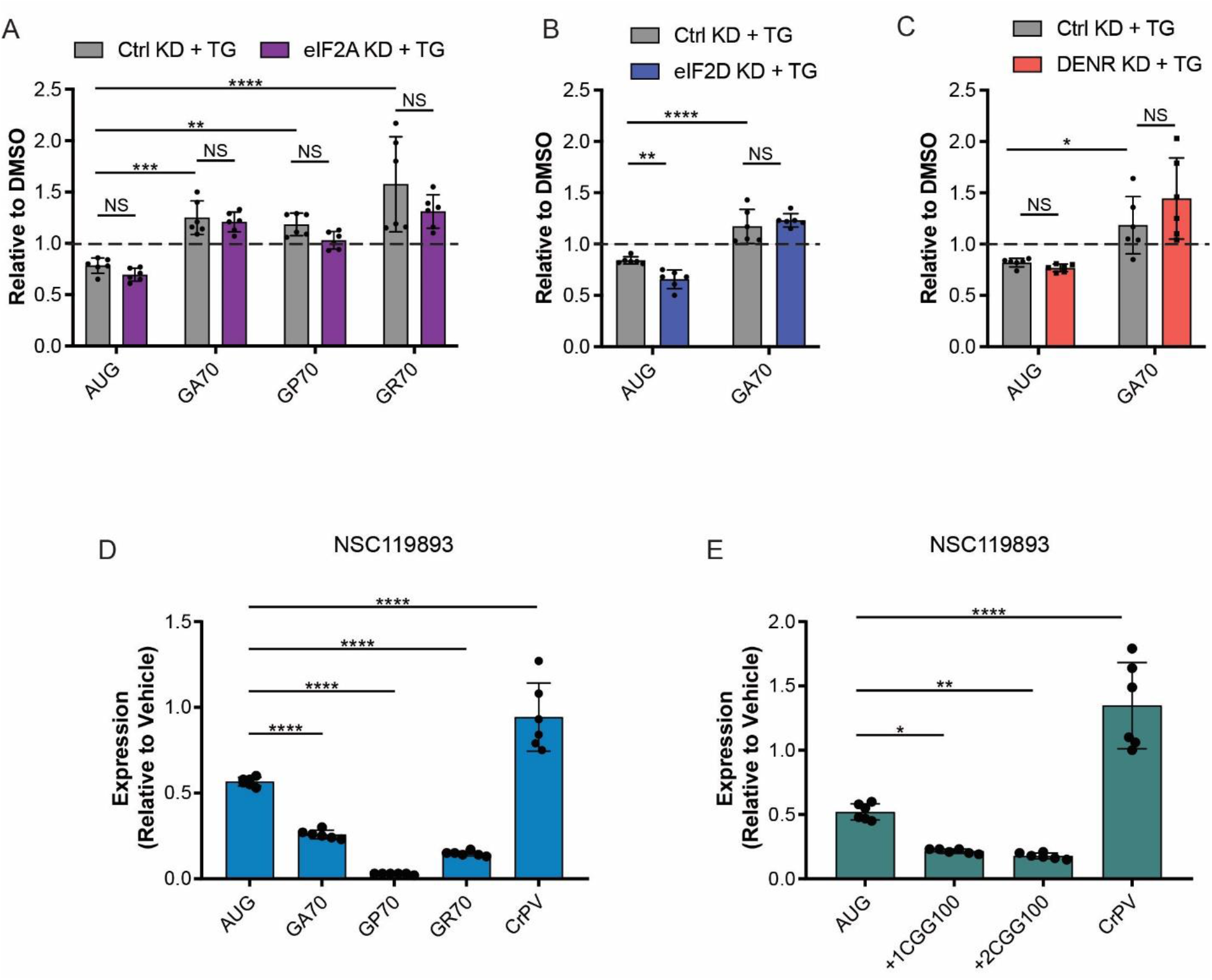
eIF2A, eIF2D, and DENR are not required for stress induced RAN translation. (A-C) HEK293 cells were transfected with eIF2A, eIF2D, or DENR siRNAs, respectively, as well as a non-targeting control siRNA for 24 hours before transfection with NLuc reporter plasmids. 19 hours post reporter transfection, HEK293 cells were treated with 2 µM thapsigargin (TG) for 5 hours, and then NLuc levels were measured. NLuc levels are expressed relative to vehicle (DMSO) treated cells. (D-E) NLuc expression for indicated reporter mRNAs expressed in rabbit reticulocyte lysate treated with 10 µM NSC119893, expressed relative to vehicle treated controls. n=6. All graphs represent mean with errors +/- standard deviation. (A-C) Two-way ANOVA with Sidak’s multiple comparison test, (D-E) One-way ANOVA with Dunnett’s multiple comparison test, * p< 0.05, ** p< 0.01, **** p< 0.0001

These results suggest that despite limited levels of functional eIF2·GTP·Met-tRNA_i_^Met^ TC, RAN translation may still utilize this factor for initiation during the ISR. Alternatively, stress-resistant RAN translation may receive an initiator tRNA through a mechanism independent of eIF2 and the additional factors assessed here. To help distinguish between these possibilities, we expressed our C9 and CGG RAN translation reporter mRNAs in a rabbit reticulocyte lysate (RRL) in the presence or absence of the small molecule NSC119893, which selectively inhibits formation of the eIF2·GTP·Met-tRNA_i_^Met^ ternary complex (33). As expected, the addition of 25 µM NSC119893 to RRL reactions greatly inhibited AUG-NLuc expression but did not affect expression of an A-capped CrPV-NLuc reporter (34) that initiates translation entirely through a non-methionyl IRES-mediated mechanism (**Figure 4D & E**). Interestingly, expression of all C9 and CGG RAN translation reporters in RRL was inhibited to an even greater extent than AUG-NLuc by NSC119893 treatment (**Figure 4D & E**). This suggests that the majority of RAN translation initiation of these repeats, under basal conditions, requires the canonical eIF2·GTP·Met-tRNA_i_^Met^ ternary complex.

### DENR KD prolongs lifespan of Drosophila expressing GGGGCC repeat

To assess the role of these factors in an *in vivo* model of repeat toxicity, we used transgenic *Drosophila* lines conditionally expressing 28 GGGGCC repeats under the UAS promoter, with the repeat inserted into either the second chromosome or third chromosome (35). When expressed with a non-targeting control shRNA in the fly eye using the eye-specific GMR-Gal4 driver, the repeat causes a severe rough eye phenotype with noticeable eye shrinkage (**Figure 5A & B**) (35).

**Figure 5:**
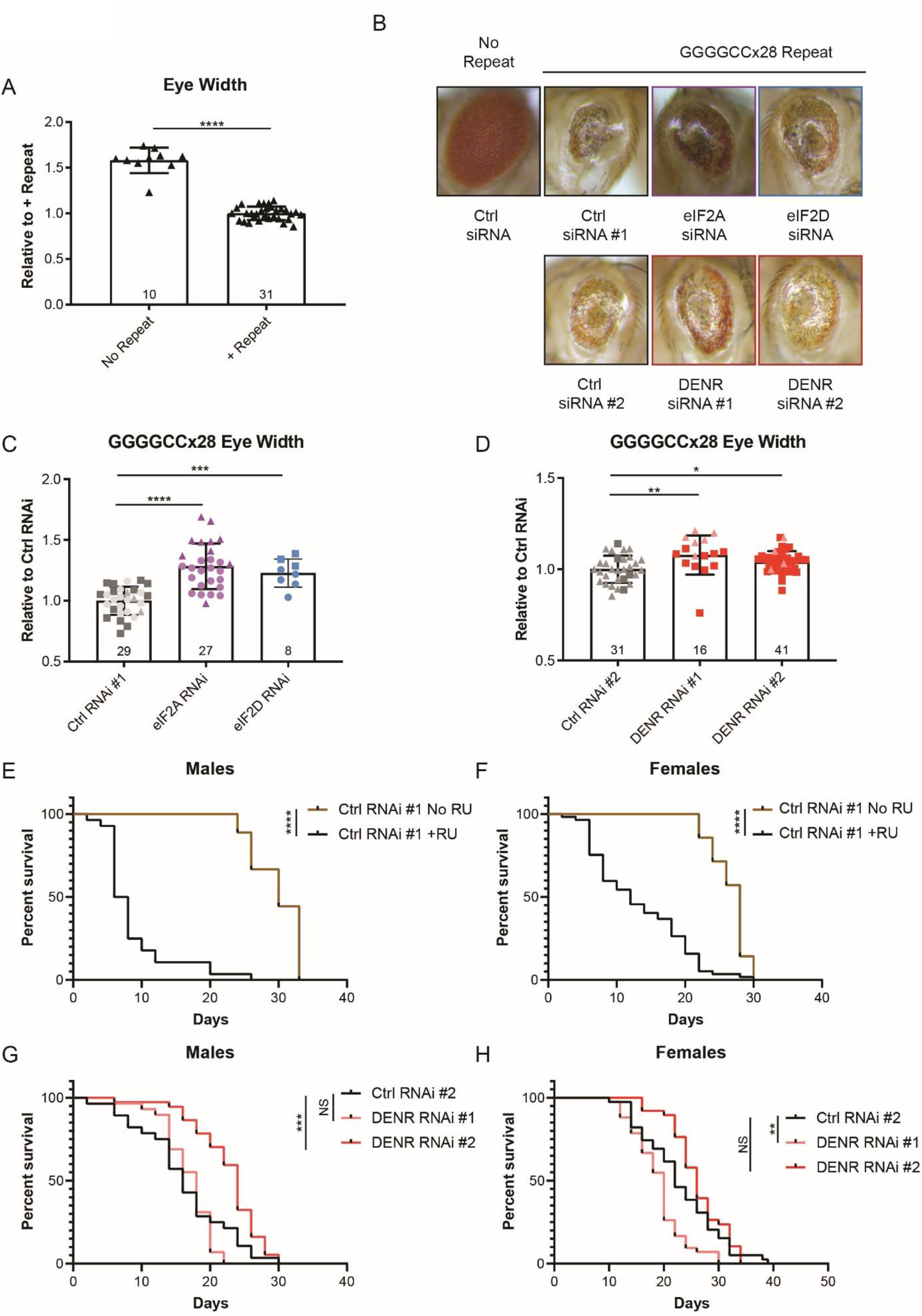
Knockdown of DENR suppresses GGGGCC associated toxicity in *Drosophila*. (A) Quantification of eye width in flies expressing control (ctrl) shRNA #1 in the absence or presence of the GGGGCCx28 repeat, under the GMR-Gal4 driver. Experimental numbers are indicated within bars, and each fly is represented by a single data point. (B) Representative images of fly eyes expressing control or CG7414 (“eIF2A”), eIF2D, or DENR shRNAs in the presence of the GGGGCCx28 repeat. (C-D) Comparison of eye width in flies expressing indicated shRNAs in the presence of the GGGGCCx28 repeat. Experimental numbers are indicated within bars, and each fly is represented by a single data point. Shape of data points represent experimental trials that data points were collected from. (E-F) Survival curve of control male and female Tub5-GS-Gal4 flies, respectively. Survival of flies with induced expression of the GGGGCCx28 repeat and control shRNA through RU486 treatment (+RU, male n = 28, female n = 57), is compared to survival of flies without induced expression (No RU, male n = 9, female n = 14). (G-H) Survival curve of RU486-treated GGGGCCx28 male and female Tub5-GS-Gal4 flies, respectively, expressing ctrl (male n = 28, female n = 39) or DENR targeting shRNA #1 (male n = 29, female n = 42) and shRNA #2 (male n = 37, female n = 38). Bar graphs represent mean with errors +/- standard deviation, * p< 0.05, ** p< 0.01, *** p< 0.001, **** p< 0.0001 by one-way ANOVA with Dunnett’s multiple comparison test. For survival curves, ** p< 0.01, *** p< 0.001, **** p< 0.0001, mantel-cox test.

We next co-expressed the GGGGCC repeat with shRNAs targeting the predicted *Drosophila* homolog of eIF2A (CG7414), eIF2D, and DENR using the GMR-Gal4 driver, and imaged the fly eyes 1-3 days post eclosion (**Figure 5B**). Using ImageJ, we measured the width of repeat-expressing fly eyes in the presence of the control versus targeting shRNAs. RNAi against eIF2A, eIF2D and DENR all modestly improved the rough of phenotype in flies maintained at 29°C (**Figure 5C & D**).

We next expressed the GGGGCCx28 repeat along with a non-targeting control shRNA ubiquitously throughout the fly using the RU-486 inducible tubulin driver, Tub5-GS Gal4. This results in a substantial decrease in the lifespan of both male and female flies (**Figure 5E & F**). We then tracked the survival of the GGGGCCx28 flies ubiquitously expressing shRNAs against eIF2A (CG7414), eIF2D, DENR, and MCTS1 compared to control shRNAs (Average KD efficiencies in **Table 3**). In doing so, we found that one shRNA targeting DENR significantly prolonged male fly survival in the presence of the repeat, although its benefit was not observed in female flies (**Figure 5G & H** and **Supplemental Fig. 4A-F**). Together these data suggest that modulating levels of non-canonical initiation factors can reduce toxicity caused by GGGGCC repeat expression in model organisms.

## Discussion

Characterizing ways in which RAN translation differs mechanistically from canonical translation has great potential for uncovering new strategies for therapeutic targeting. Previous work from our group and others revealed the ability of C9 and CGG RAN translation to continue during ISR activation, a condition that impairs canonical translation (23-26). As the ISR inhibits functional eIF2 ternary complex formation, we hypothesized that RAN translation uses factors that can function in place of eIF2. We thus investigated the role of eIF2A, eIF2D, and DENR/MCTS1 in supporting C9 and CGG RAN translation. While our results indicate that these factors are not responsible for promoting increased RAN translation during the ISR, they do support a role for non-canonical initiation factors, particularly the DENR/MCTS1 complex, in RAN translation initiation.

Despite being first identified nearly 50 years ago (8, 9), eIF2A is still a poorly understood translation factor. Recent work has established it as a modifier of both C9RAN translation and RAN translation of CCUG and CAGG repeats causative of myotonic dystrophy type 2 (25, 30). While we found that deletion of eIF2A more greatly impaired RAN translation than AUG-initiated translation in *in vitro* lysates, we did not see a robust or consistent inhibition following eIF2A KD in HEK293 cells. One possible explanation for the lack of effect in a cell-based system is that the interplay between eIF2A’s role in regulating translation during cellular stress (7, 31, 32), the cellular stress resulting from transient transfections, and the increase in RAN translation during cellular stress conditions (23-26), could obscure the role eIF2A plays in regulating RAN translation under the non-stressed conditions in *in vitro* translation lysates. In the context of previously published work (25, 30), this suggests that eIF2A function may be highly context dependent, with the factors that regulate its activity still unknown. This is consistent with conflicting reports on whether or not eIF2A is involved in IRES-mediated translation of the hepatitis C virus mRNA (28, 31, 36).

Unlike eIF2A KD, KD of the DENR/MCTS1 complex, through use of either DENR or MCTS1 targeting siRNAs, greatly reduced RAN translation across all three reading frames of GGGGCCx70 reporters, and in both the +1 and +2 reading frames of CGGx100 reporters. This inhibition was specific to RAN translation, as DENR/MCTS1 KD had little effect on near-AUG reporters lacking expanded repeats, or on AUG-initiated repeat-containing reporters. The dependence on both non-AUG initiation and the expanded repeat is intriguing. While the DENR/MCTS1 complex can support translation initiation through delivery of initiator tRNAs to the ribosome (16), its main described functions to date are in ribosome recycling and re-initiation (13-15). As translation through GC rich repeats may trigger ribosome stalling where choices must be made between re-initiation and ribosomal recycling pathways, RAN translation may uniquely require the combination of non-canonical initiation and assistance with translational elongation.

Unfortunately, unlike eIF2A, the DENR/MCTS1 complex is essential for life (15). While the conditions of our knockdown do not impair global translation as measured by polysome profiling and our AUG-NLuc reporters, they do reduce FFLuc reporter expression, which is less stable and may be an indication of reduced cellular fitness. Future work aimed at better understanding the mechanism by which DENR/MCTS1 supports translation may reveal ways to specifically target RAN translation without interfering with DENR/MCTS1’s essential functions.

Our work in *Drosophila* demonstrated partial rescue associated with knockdown of either eIF2A or DENR expression, with the effects of DENR being more robust. While these findings are promising and provide evidence for DENR KD mediated effects on repeat associated toxicity across multiple assays, these models lack the normal genomic sequence context for the C9orf72 repeat, which places limitations on its transferability into more complex organisms or human patients. Future studies will be needed to understand the strength of its selective effects in patient derived cells with full-length repeats in their endogenous sequence context.

In sum, we have studied the role of three alternative ternary complexes for their ability to selectively modulate RAN translation at two different repeat elements and in a *Drosophila* model of C9 ALS/FTD. These studies suggest that the DENR/MCTS1 complex in particular has roles in RAN translation and is worthy of further study in human neuronal model systems.

## Experimental methods

### RNA synthesis

RNAs were *in vitro* transcribed from PspOMI-linearized pcDNA3.1(+) reporter plasmids using HiScribe T7 High Yield RNA Synthesis Kit (NEB) with 3’-O-Me-m^7^GpppG anti-reverse cap analog (ARCA) or ApppG cap (NEB) added at 8:1 to GTP for a capping efficiency of ∼90%, as previously described (23, 27). After RNA synthesis, DNA templates were removed with RNase-free DNaseI (NEB), and RNAs were poly-adenylated with *E. coli* Poly-A Polymerase (NEB) as previously described (23, 27). mRNAs were then clean and concentrated with RNA Clean and Concentrator-25 Kit from Zymo Research and run on a denaturing formaldehyde RNA gel to verify mRNA size and integrity. All reporter sequences have been previously published (23, 27).

### Rabbit reticulocyte lysate in vitro translation

mRNAs were *in vitro* translated with Flexi Rabbit Reticulocyte Lysate System (Promega), as previously described (23, 27). 10 µL reactions for luminescence assays were programmed with 3 nM mRNA and contained 30% RRL, 10 µM amino acid mix minus methionine, 10 µM amino acid mix minus leucine, 0.5 mM MgOAc, 100 mM KCl, and 0.8 U/µL Murine RNAse Inhibitor (NEB). 25 µM NSC119893 (the Chemical Repository at the National Cancer Institute’s Developmental Therapeutics Program) or equal volume DMSO was added to the RRL reaction mix to constitute 1% of final reaction volume (0.1 µL/reaction). Reactions were incubated at 30°C for 30 minutes before termination by incubation at 4°C. Samples were then diluted 1:7 in Glo Lysis Buffer (Promega) and incubated 1:1 for 5 minutes in the dark in opaque 96 well plates with NanoGlo Substrate freshly diluted 1:50 in NanoGlo Buffer (Promega). Luminescence was measured on a GloMax 96 Microplate Luminometer.

### Cell line information and maintenance

HEK293 cells were purchased from American Type Culture Collection (ATCC CRL-1573). They were maintained within 37°C incubators at 5% CO_2_ in DMEM + high glucose (GE Healthcare Bio-Science, SH30022FS) supplemented with 9.09% fetal bovine serum (50 mL added to 500 mL DMEM; Bio-Techne, S11150).

eIF2A KO (Cat # HZGHC002650c001) and isogenic control HAP1 cells were purchased from Horizon Discovery (Cambridge, UK) (28, 29). They were maintained within 37°C incubators at 5% CO_2_ in IMDM (Invitrogen, 124400-61) supplemented with 9.09% fetal bovine serum (50 mL added to 500 mL IMDM; Bio-Techne, S11150).

### HAP1 cell lysate in vitro translation

Lysates were prepared using a previously developed protocol within our lab (23). HAP1 cells were trypsinized, centrifuged at 200xg for 5 minutes, and the cell pellet washed once with 1X PBS. Cell pellets were then weighed and resuspended in RNAse-free hypotonic buffer containing 10 mM HEPES-KOH (pH 7.6), 10 mM potassium acetate, 0.5 mM magnesium acetate, 5 mM DTT, and EDTA-free protease inhibitor cocktail (Roche), 250 µL buffer added per 200 mg cells (23). Resuspended cells were incubated on ice for 20 minutes, passed 10X through a 27G syringe, incubated for another 20 minutes on ice, and centrifuged at 10,000xg for 10 minutes at 4°C to pellet cell debris. The supernatant was recovered, and total protein quantified with a BCA assay. Lysates were diluted to 8 μg/μl protein in the hypotonic lysis buffer and stored in single use aliquots at -80°C.

For *in vitro* translation reactions, 8 µg lysate was supplemented to final concentrations of 20 mM HEPES-KOH (pH 7.6), 44 mM potassium acetate, 2.2 mM magnesium acetate, 2 mM DTT, 20 mM creatine phosphate (Roche), 0.1 µg/µl creatine kinase (Roche), 0.1 mM spermidine, and on average 0.1 mM of each amino acid (23). *In vitro* transcribed reporter mRNAs were added to 4 nM. Translation assays and luminescence measurements were then performed as with RRL reactions.

### Cell transfection, drug treatments, and analysis

For luminescence assays, HEK293 cells were seeded in 96-well plates and transfected 24 hours later at 40-50% confluency with siRNAs using Lipofectamine RNAiMax. The following amount of stealth siRNA (Thermo) was added per well, to achieve maximum knockdown efficiency: 0.1 pmol eIF2A siRNA, 1 pmol DENR siRNA, 2 pmol MCTS1 siRNA, 4 pmol eIF2D siRNA. The control non-targeting siRNA was added at same concentration as the targeting siRNA it was compared to. siRNA sequences are listed in **Table 1**.

**Table 1.**
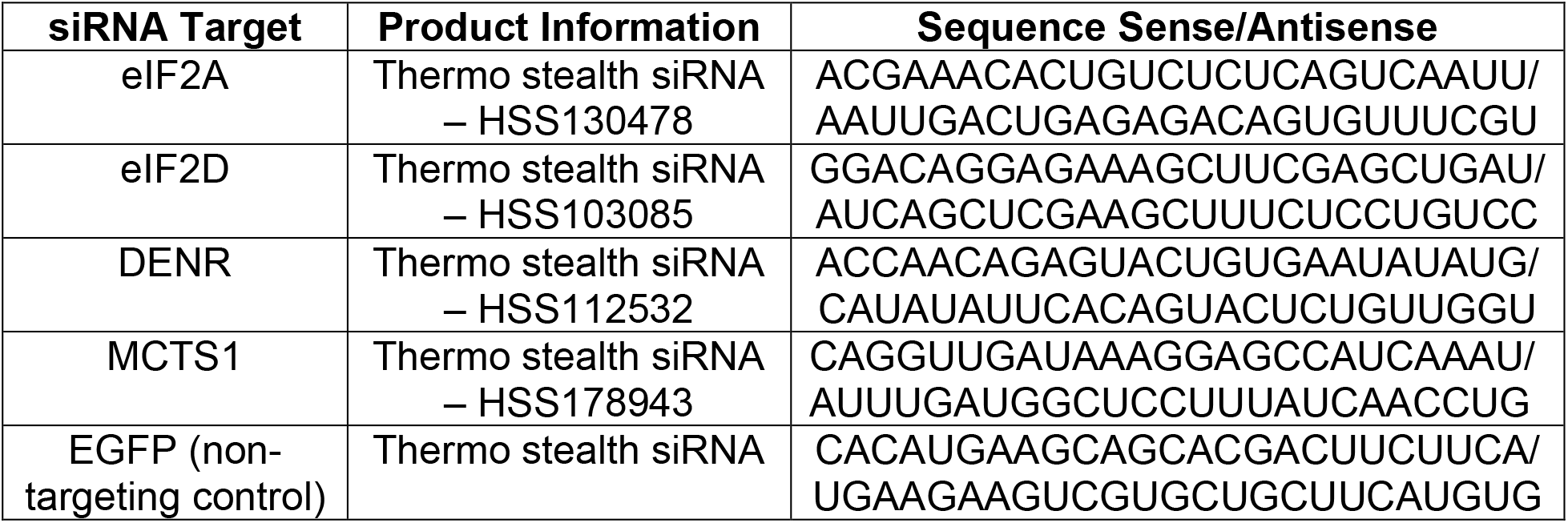
siRNA product information

Reporters were transfected into cells ∼24 hours post siRNA transfections, when cells were 80-90% confluent. DNA transfections were performed in triplicate with FuGene HD at a 3:1 ratio to DNA, with 50 ng NLuc reporter DNA and 50 ng pGL4.13 FLuc reporter added per well. RNA transfections were performed with *Trans*IT-mRNA Transfection Kit from Mirus Bio, per manufacturer’s recommended protocol, with 90 ng reporter mRNA and 200 ng pGL4.13 FLuc control DNA added to each well in triplicate.

For TG experiments, 2 µM TG or equal volume DMSO was added to HEK293 cells 19 hours post transfection, for 5 hours.

Cells were lysed 24 hours post transfection with 60 μL Glo Lysis Buffer for 5 minutes at room temperature. 25 μL of lysate was mixed with NanoGlo Substrate prepared as for RRL reactions, and 25 μL of ONE-Glo Luciferase Assay System (Promega), for 5 minutes in the dark, in opaque 96-well plates. Luminescence measurements were obtained as with RRL reactions.

For western blots, HEK293 cells were seeded in 12-well plates and transfected 24 hours later, at 40-50% confluency, with siRNAs using RNAiMax. Where indicated, 24 hours post transfections, at 80-90% confluency cells, were transfected with 500 ng NLuc reporter DNAs and 4:1 FuGene HD. 48 hours post siRNA transfection, cells were lysed in 300 μL RIPA buffer with protease inhibitor for 30 minutes at 4°C. Lysates were homogenized by passing through a 28G syringe, mixed with 6X sample buffer, and stored at -20°C.

### Western blots

All samples for western blot were run on 12% SDS-polyacrylamide gels at 150V for ∼90 minutes. For western blot analysis of HAP1 lysates, 60 µg protein was loaded per well. For HEK293 western blots, 30 µL lysate for each sample was loaded. Gels were transferred to PVDF membranes either overnight at 30V and 4°C, or for 2.5 hours at 320 mAmps and 4°C. Membranes were blocked with 5% non-fat dairy milk, and all antibodies were diluted in 5% non-fat dairy milk. Primary antibodies information and probing conditions are listed in **Table 2**. For eIF2A, eIF2D, DENR, MCTS1, and FLAG western blots, HRP secondary antibodies were applied at 1:10,000, with 1-hour incubations at room temperature. Bands were then visualized on film. For GAPDH loading controls, LiCor IRDye secondary antibodies were applied at 1:10,000, with 1-hour incubations at room temperature, and bands visualized with LiCor Odyssey CLx Imaging Systems.

**Table 2.** Primary antibody information

| **Antibody Target** | **Product Information** | **Species** | **Probing Conditions** |
| --- | --- | --- | --- |
| eIF2A | Protein Tech, 11233-1-AP | Rabbit | 1:8000 - 1 hour at room temperature |
| eIF2A | Abcam; ab169528 | Rabbit | 1:1000 – overnight at 4C |
| eIF2D | Protein Tech, 12840-1-AP | Rabbit | 1:1000 – overnight at 4C |
| DENR | Sigma, Clone 1H3,<br>WH0008562M1 | Mouse | 1:1000 – overnight at 4C |
| MCTS1 | Sigma, SAB2701331<br>(discontinued) | Rabbit | 1:500 – overnight at 4C |
| FLAG | Sigma, M2, F1804 | Mouse | 1:1000 – overnight at 4C |
| GAPDH | SCBT, sc-32233 | Mouse | 1:1000 – 1 hour at room temperature |

### Polysome profiling

24-hour post indicated knockdown, HEK293 cells were treated with 100 μg/ml cycloheximide (CHX) for 5 minutes at 37°C. They were then harvested as previously described (37). Briefly, they were transferred to ice and washed with 5.0 mL ice-cold PBS containing 100 μg/ml CHX, collected by scraping in cold PBS+CHX, and pelleted at 234xg and 4°C for 5 minutes. PBS was aspirated and pellets re-suspended in polysome-profiling lysis buffer (20mM Tris–HCl (pH 7.5), 150mM NaCl, 15mM MgCl2, 8% (vol/vol) glycerol, 20U/ml SUPERase, 80U/ml murine RNase inhibitor, 0.1mg/ml heparin, 100μg/ml CHX, 1mM DTT, 1× EDTA-free protease inhibitor cocktail, 20U/ml Turbo DNase, 1% Triton X-100) (37). Lysates were passed through a 20G needle 10x and incubated on ice for 5 minutes. Cellular debris was pelleted at 14,000xg and 4°C for 5 minutes, and supernatant transferred to a fresh tube. Total lysate RNA was estimated by NanoDrop. Lysates were flash-frozen in liquid nitrogen and stored at -80°C until fractionation.

Sucrose gradients were prepared by successively freezing equal volumes of 50, 36.7, 23.3, and 10% sucrose (wt/vol) in 12-ml Seton tubes. Sucrose-gradient buffer consisted of 20mM Tris–HCl (pH 7.5), 150mM NaCl, 15mM MgCl2, 10U/ml SUPERase, 20U/ml murine RNase inhibitor, 100μg/ml CHX, and 1mM DTT (37). Prior to use, gradients were allowed to thaw and linearize overnight at 4°C. For fractionation, approximately 250 μg total RNA was applied to the top of the sucrose gradient. Gradients were spun at 151,263xg and 4°C for 3 hours using a Beckman Coulter Optima L-90K ultracentrifuge and SW 41 Ti swinging-bucket rotor. Gradients were fractionated with Brandel’s Gradient Fractionation System, measuring absorbance at 254 nm. The detector was baselined with 60% sucrose chase solution, and its sensitivity set to 1.0. For fractionation, 60% sucrose was pumped at a rate of 1.5 mL/min. Brandel’s PeakChart software was used to collect profile data.

### Drosophila work

The GGGGCCx28 repeat, along with 30 nt on both ends in the first intron of the gene C9ORF72 were PCR cloned from the genomic DNA from fibroblasts of an ALS patient of Central Biorepository of University of Michigan, and placed to the 5′ upstream in the +1 reading frame (GP) relative to the GFP gene in the vector PGFPN1 (Clonetech). The repeat and GFP were then subcloned into the NotI site of vector pUAST and the sequence was verified for the repeat length and relative reading frame. This vector was used to generate transgenic flies by standard p-element insertion (Best Gene, CA) (35).

Information on the GAL4 and shRNA lines interrogated is included in **Table 3**. For eye shrinkage experiments, male flies containing the UAS-GGGGCCx28 and a UAS-shRNA transgene were crossed to GMR-GAL4 virgin females at 29°C. One eye of each resulting progeny was imaged 0-2 days post eclosion using Leica M125 stereomicroscope and a Leica DFC425 digital camera. Eye widths were then measured with ImageJ, and normalized to values obtained to flies expressing control shRNAs.

**Table 3.**
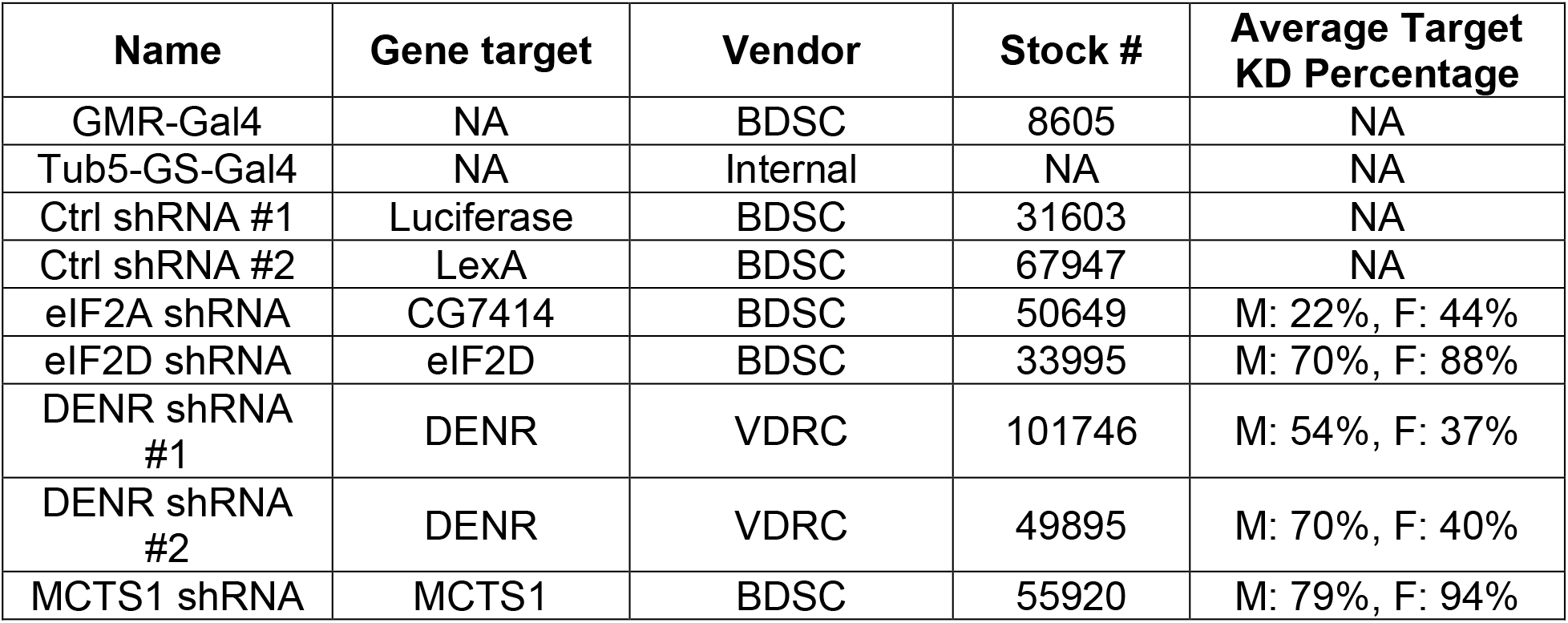
*Drosophila* line information

For survival experiments, male flies containing the UAS-GGGGCCx28 and a UAS-shRNA transgene were crossed to Tub5-GAL4 GeneSwitch (GS) virgin females at 25°C. 0-2 days post eclosion, resulting progeny were placed on SY10 food containing 200 µM RU486. Male and female progeny were housed separately, with no more than 28 flies per tube. RU486 food was changed every 2 days, with number of dead flies counted during each flip.

For real time quantitative PCR, total RNA was isolated from 19-26 flies per sex per genotype with TRIzol reagent (Invitrogen). 10 µg of isolated RNA was treated with TURBO DNase (Ambion) to remove contaminating DNA and was clean and concentrated with RNA Clean and Concentrator-25 Kit (Zymo Research). 1 µg of cleaned RNA was used to generate cDNA with iScript reverse transcriptase per the manufacturer’s protocol (Bio-Rad). 20 µL quantitative real-time PCR reactions with 100 ng of cDNA input were carried out using TaqMan Fast Advance Master Mix (Applied Biosystems, 4444557) on an Applied Biosystems Quant Studio 3 machine for 40 cycles using fast cycling parameters (95C for 20s, 95C for 1s, 60C for 20s). All runs included a standard dilution curve representing 2x to 0.02x of the RNA concentration utilized for all primer sets to ensure linearity. Equivalent efficiency of individual primer sets was confirmed prior to data analysis. All samples were run in triplicate. The level of mRNA of interest was normalized to RPL32 mRNA and expressed as the change in gene expression to control lines.

## Acknowledgements

We thank all members of the Todd Lab for their input and assistance on this project. In particular, we thank Udit Sheth, Kristina Zheng, Katyanne Calleja, and Sofia Bennetts for assistance maintaining fly lines used in these studies. We also thank the Pletcher Lab at the University of Michigan for making our fly food.

This work was funded by NIH grants P50HD104463, R01NS099280 and R01NS086810 to PKT, and F31NS100302 to KMG. PKT was also supported by VA grant BLRD BX004842, and by private philanthropic support. KMG and SLM were additionally supported by the Cellular and Molecular Biology Graduate Program at the University of Michigan, T32GM007315. IM was supported by an Alzheimer’s Association Research Fellowship.

## Supplemental Figures

**Figure S1:**
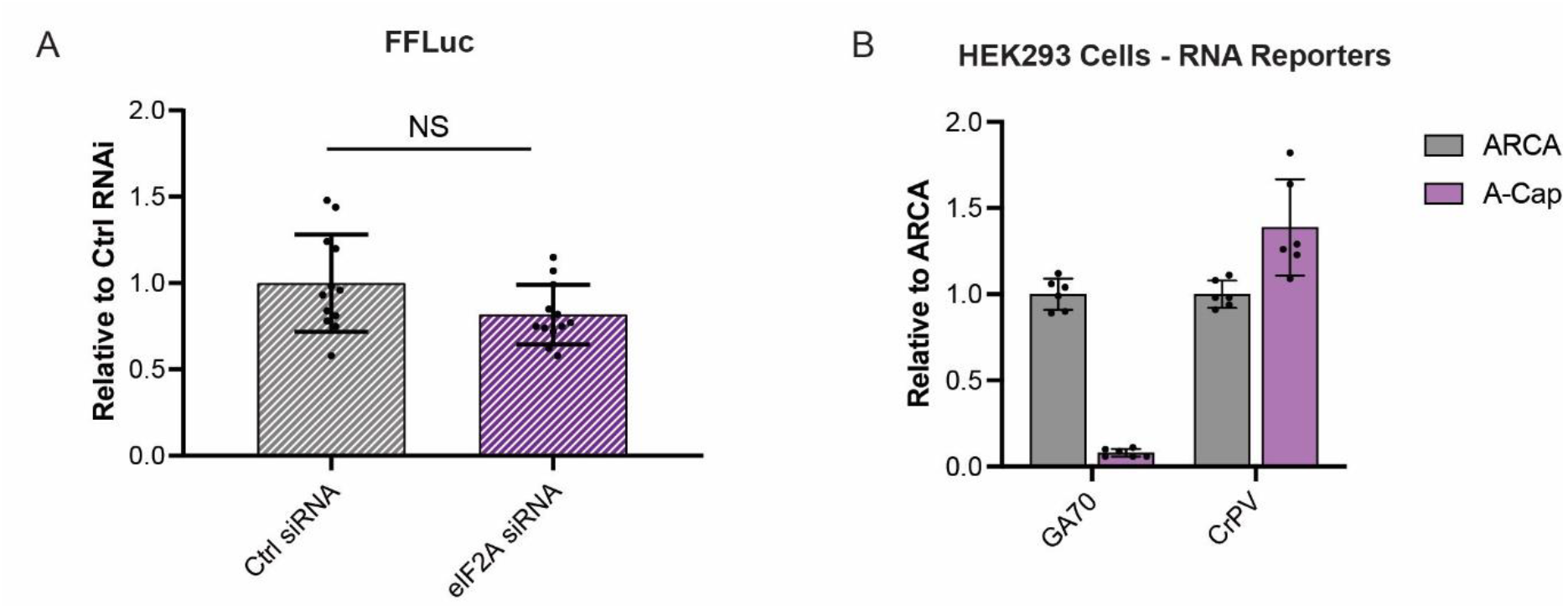
Effect of eIF2A knockdown and 5′ m7G cap on translation. (A) Expression of FFLuc reporter in wells co-transfected with AUG-NLuc, 24 hours post transfection with non-targeting or eIF2A siRNAs. FFLuc levels are expressed relative to levels in cells transfected with the non-targeting siRNA, n=12. (B) NLuc expression of indicated A-capped reporter mRNAs expressed relative to corresponding ARCA reporter mRNAs, transfected into HEK293 cells 24 hours post transfection with a non-targeting siRNA, n=6. All graphs represent mean with errors +/- standard deviation.

**Figure S2:**
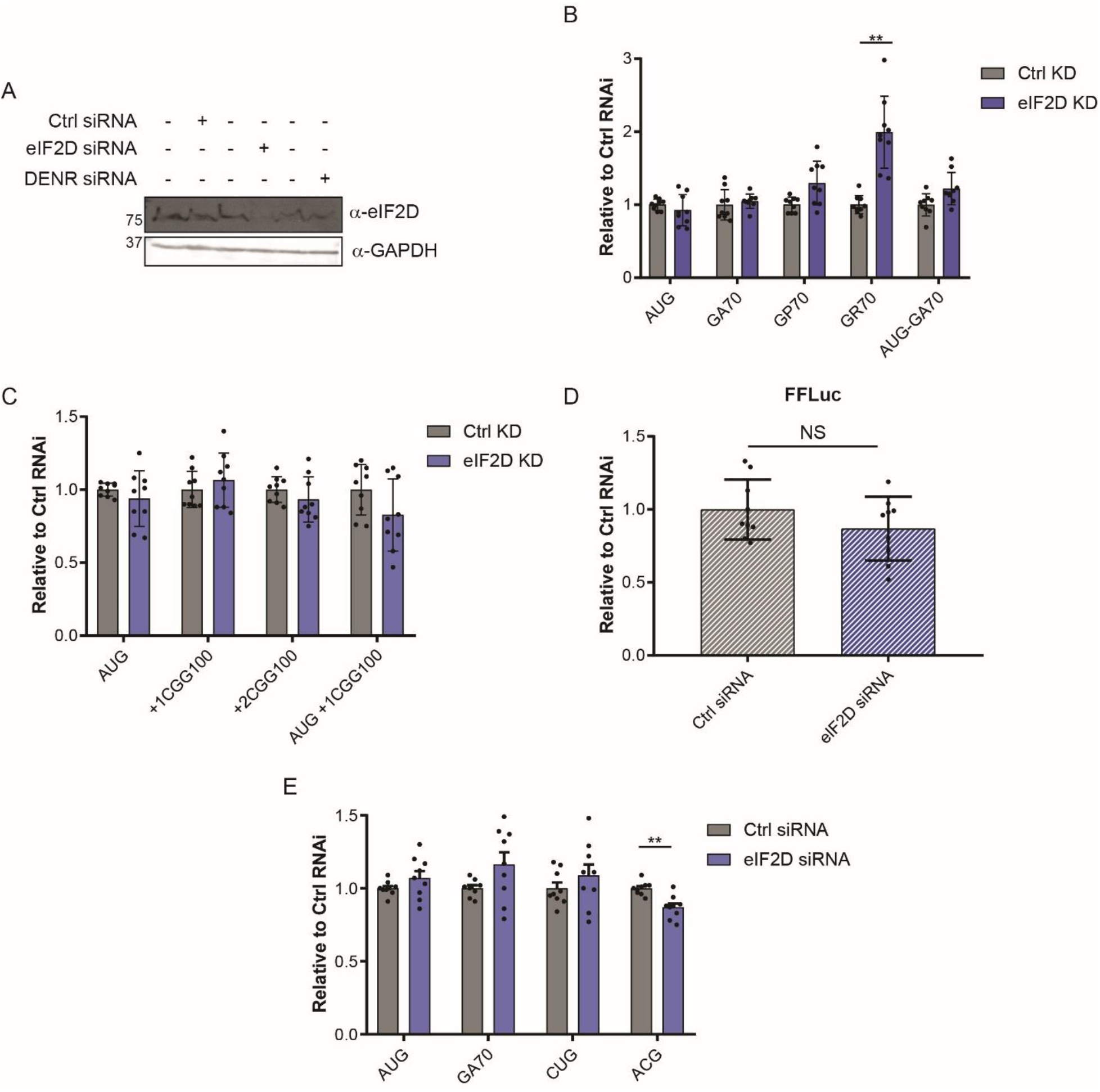
Knockdown of eIF2D does not alter RAN translation reporter expression. (A) Western blot showing efficiency of eIF2D KD in HEK293 cells 48 hours post transfection with an eIF2D targeting siRNA. GAPDH was used as a loading control. (B-C) NLuc expression of indicated reporters expressed from DNA plasmids transfected into HEK293 cells 24 hours post transfection with non-targeting or eIF2D siRNAs. NLuc levels are expressed relative to levels in cells transfected with the non-targeting siRNA, n=9. (D) Expression of FFLuc reporter in wells co-transfected with AUG-NLuc, 24 hours post transfection with non-targeting or eIF2D siRNAs. FFLuc levels are expressed relative to levels in cells transfected with the non-targeting siRNA, n = 9. (E) NLuc expression of indicated reporters expressed from DNA plasmids transfected into HEK293 cells 24 hours post transfection with non-targeting or eIF2D siRNAs. NLuc levels are expressed relative to levels in cells transfected with the non-targeting siRNA, n=9. All graphs represent mean with errors +/- standard deviation, ** p< 0.01 (B-C, E) Two-way ANOVA with Sidak’s multiple comparison test, (F) two-tailed unpaired t-test.

**Figure S3:**
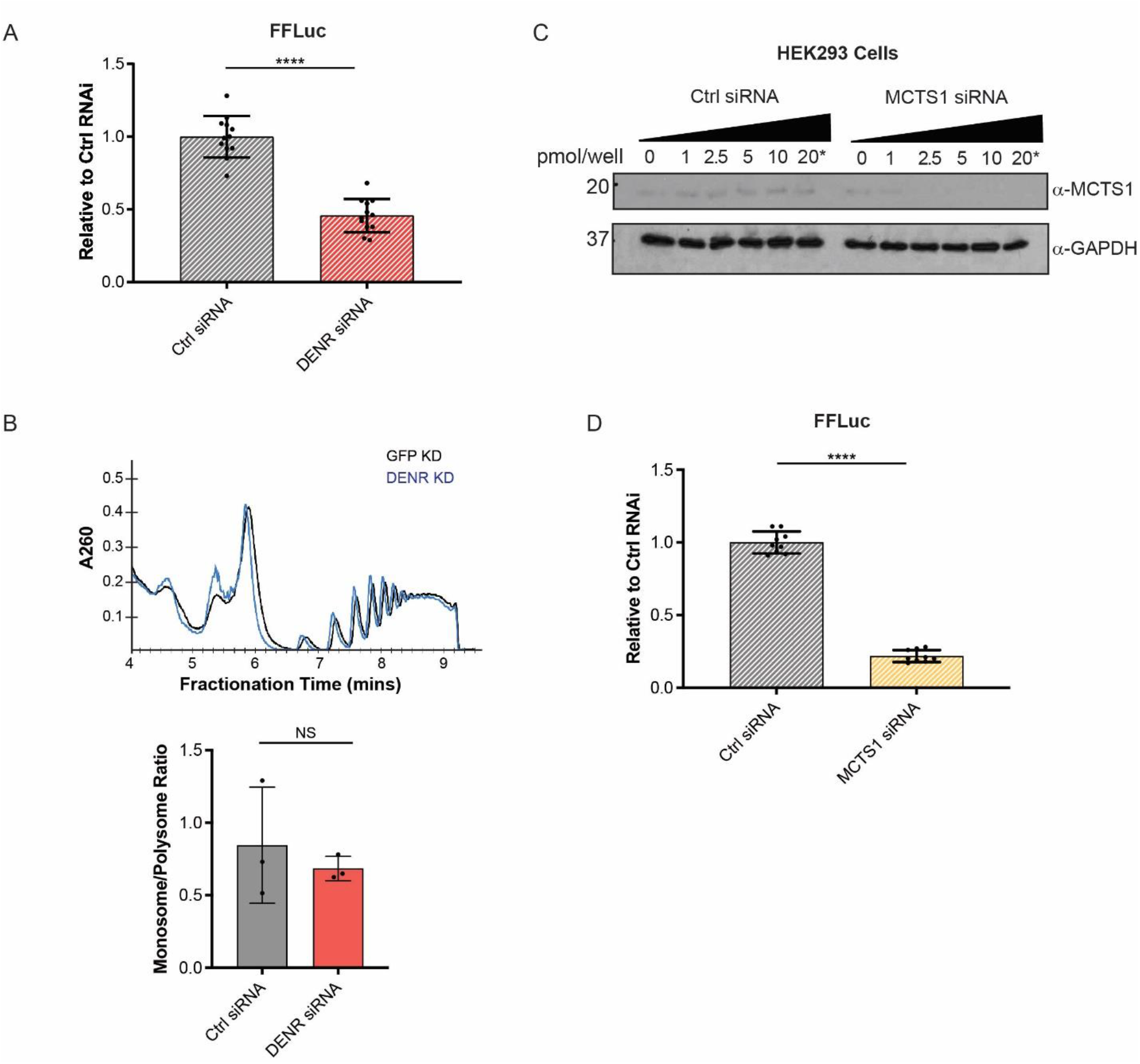
Effects of DENR and MCTS1 knockdown on canonical translation. (A) Expression of FFLuc reporter in wells co-transfected with AUG-NLuc, 24 hours post transfection with non-targeting or DENR siRNAs. FFLuc levels are expressed relative to levels in cells transfected with the non-targeting siRNA, n=12. (B) Representative polysome profiles and monosome/polysome ratios of HEK293 cells 24 hours post transfection with non-targeting or DENR siRNAs, n=4. (C) Western blot showing efficiency of MCTS1 KD in HEK293 cells following increasing concentrations of MCTS1 siRNA. The starred lane indicates siRNA concentration used in subsequent experiments. GAPDH was used as a loading control. (D) Expression of FFLuc reporter in wells co-transfected with AUG-NLuc, 24 hours post transfection with non-targeting or MCTS1 siRNAs. FFLuc levels are expressed relative to levels in cells transfected with the non-targeting siRNA, n=9. All graphs represent mean with errors +/- standard deviation, *** p< 0.001, **** p< 0.0001, two-tailed unpaired t-test.

**Figure S4:**
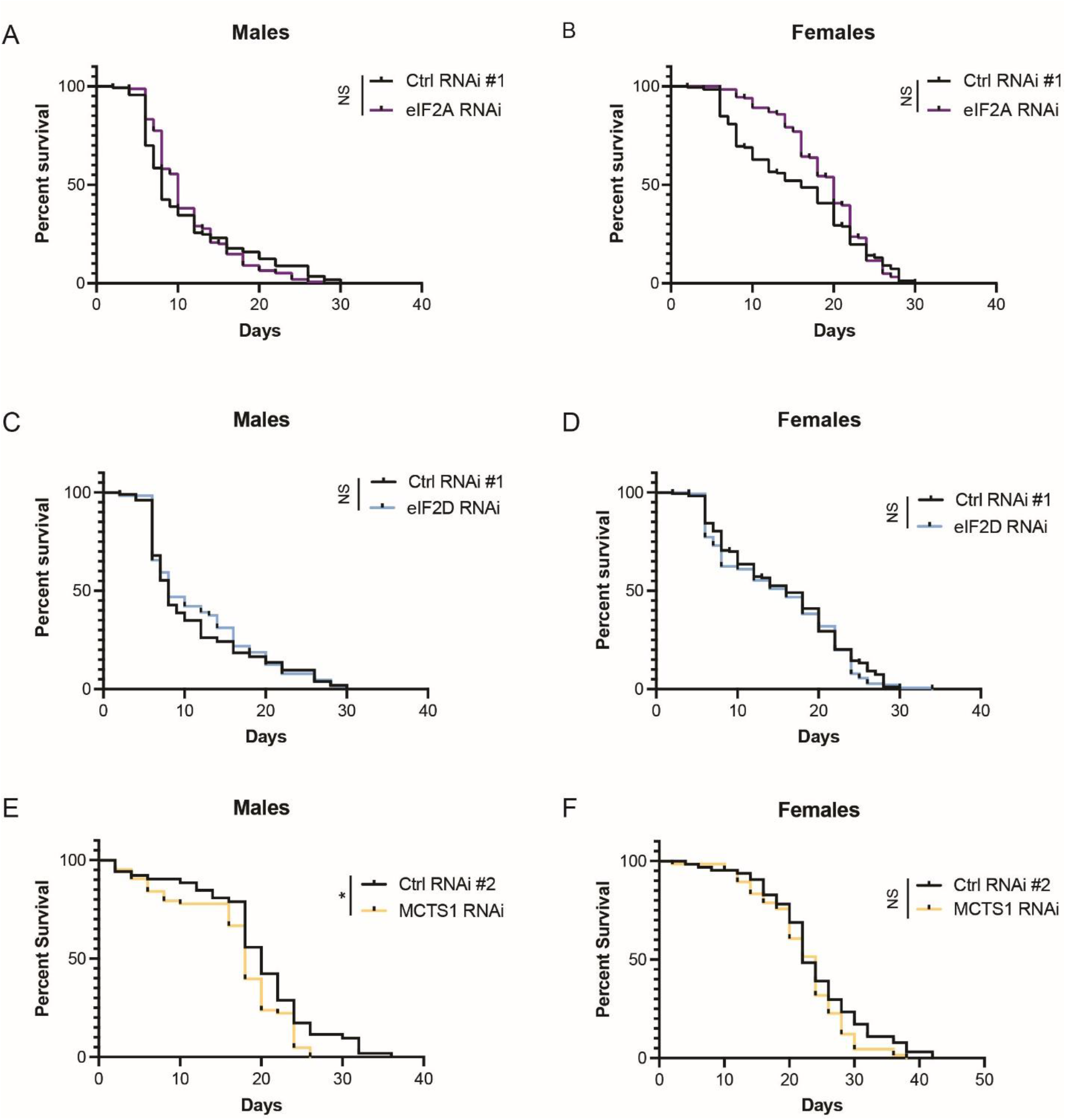
Effects of eIF2A, eIF2D and MCTS1 knockdown on GGGGCC repeat-suppressed *Drosophila* survival. (A-F) Survival curves of RU486-treated GGGGCCx28 male and female Tub5-GS-Gal4 flies, expressing ctrl or eIF2A, eIF2D, or MCTS1 targeting shRNAs. (A) Ctrl n = 113, eIF2A n = 155. (B) Ctrl n = 177, eIF2A n = 182. (C) Ctrl n = 103, eIF2D n = 63. (D) Ctrl n = 173, eIF2D n = 141. (E) Ctrl n = 52, MCTS1 n = 63. (F) Ctrl n = 64, MCTS1 n = 66. *p< 0.05 by mantel-cox test.

